# The effect of smoking on the brain revealed by using electronic cigarettes with concurrent fMRI

**DOI:** 10.1101/107771

**Authors:** Matthew B Wall, Alexander Mentink, Georgina Lyons, Oliwia S Kowalczyk, Lysia Demetriou, Rexford D Newbould

## Abstract

Cigarette addiction is driven partly by the physiological effects of nicotine, but also by the distinctive sensory and behavioural aspects of smoking, and understanding the neural effects of such processes is vital. There are many practical difficulties associated with subjects smoking in the modern neuroscientific laboratory environment, however electronic cigarettes obviate many of these issues, and provide a close simulation of smoking tobacco cigarettes. We have examined the neural effects of ‘smoking’ electronic cigarettes with concurrent functional Magnetic Resonance Imaging (fMRI). The results demonstrate the feasibility of using these devices in the MRI environment, and show brain activation in a network of cortical (motor cortex, insula, cingulate, amygdala) and sub-cortical (putamen, thalamus, globus pallidus, cerebellum) regions. Concomitant relative deactivations were seen in the ventral striatum and orbitofrontal cortex. These results reveal the brain processes involved in (simulated) smoking for the first time, and validate a novel approach to the study of smoking, and addiction more generally.

## Introduction

Smoking is a major worldwide health problem, and current treatments for cigarette addiction are only partly effective. The behavioural and sensory aspects of smoking are thought to be important aspects of cigarette addiction, independently of the effects of nicotine (Rose, 2006; Balfour, 2004). Aspects of the multi-sensory (visual, tactile, olfactory, taste) experience of smoking can act as a powerful cue that reliably triggers craving and withdrawal symptoms in smoking addicts (Niaura et al., 1998). This is consistent with the incentive-sensitization theory of addiction (Robinson & Berridge, 1993; 2001), which proposes that neutral stimuli can promote drug-seeking behaviour through association with drug effects, and that this is mediated through sensitization of particular brain systems, principally the ventral striatum.

Research on the neural effects of smoking addiction has generally followed two largely independent paths. One has been concerned with the neurophysiological effects of nicotine and has generally used alternative routes of administration (intravenous, cutaneous patch, oral; e.g. Stein et al., 1998; Yamamoto Rohan & Goletiani, 2013; Cole et al., 2010). The second has used cigarette cues (usually images) to stimulate craving in (usually nicotine-abstinent) smokers (e.g. David et al., 2005; Janes et al., 2010). The former have provided usefully pure measures of the pharmacological effects of nicotine, and the latter have helped illuminate craving and drug-seeking mechanisms. However very few studies have investigated the neural effects of the most powerful cue associated with nicotine: the behavioural and multi-sensory repertoire of smoking itself. A small number of Positron Emission Tomography (PET) studies have involved subjects smoking while in a PET scanner. Berridge et al. (2010) used radio-labelled ^[11C]^nicotine to investigate the pharmacokinetics of nicotine absorption, and Barrett et al (2004) investigated the hedonic properties of smoking with the use of ^[11C]^raclopride to index dopamine release. The latter study showed that smoking-related changes in euphoria were related to dopamine release in the caudate and putamen, though not in the ventral striatum. Domino et al. (2013) used regular and denicotinized cigarettes to examine dopamine release; both showed effects on dopamine throughout the striatum, but the denicotinized cigarettes led to significantly less dopamine release. Cosgrove et al. (2014) also identified a potential sex difference in striatal dopamine release, with male subjects showing a more consistent and rapid response to smoking.

These PET studies are restricted, by the nature of the method, to examining a single neurotransmitter system. To date, there have been no investigations of active smoking using a more general neuroimaging method which could reveal effects across the entire brain, such as functional Magnetic Resonance Imaging (fMRI)^1^. This is likely because a host of practical, health, and safety issues largely preclude the use of traditional (i.e. combustible tobacco) cigarettes in the modern neuroscientific laboratory environment. For example: most MRI scanners are enclosed and restrictive, air capture/filtration systems would be required to deal with the smoke, and the procedure may require a naked flame. In addition local or national regulations may prohibit smoking within research institutions or hospital sites (e.g. Health Act 2006, in the United Kingdom).

Many of these issues may be obviated by the use of electronic cigarettes, or more accurately, Electronic Nicotine Delivery Systems (ENDS; Nutt et al., 2014). These are relatively novel consumer products that deliver nicotine, and are designed to provide a closer simulation of smoking tobacco cigarettes than previous nicotine-containing products. Use of ENDS does not involve combustion, and most produce relatively small amounts of vapour that evaporates within a few seconds. Evidence suggests that ENDS are effective at reducing withdrawal symptoms in nicotine-deprived smokers (Dawkins et al., 2012) and preliminary data show that ENDS may be effective at helping smokers quit, or reducing their smoking (Rahman et al., 2015; Bullen et al., 2013; although see Kalkhoran et al., 2016 for a contrary view). As such, they may hold great potential as a replacement for traditional cigarettes, and be a major benefit to public health (Nutt et al., 2014). As a novel technology, their effects on the brain, general health, and patterns of tobacco use is still largely unknown, and a recent editorial in Nature Neuroscience (2014) highlighted the urgent need for more research on ENDS.

We have explored the use of ENDS in combination with functional Magnetic Resonance Imaging (fMRI) in order to visualise brain activity related to active smoking, or more strictly, to the close simulation of active cigarette smoking that ENDS provide. To achieve this we first undertook extensive testing of commercially available ENDS to assess the devices for magnetic susceptibility and potential effects on MRI image quality. After successfully identifying a device that had minimal magnetic susceptibility and no discernible effects on image quality, 11 healthy smokers completed a scanning session that included active smoking fMRI tasks. In the first task subjects were instructed to ‘smoke’ (i.e. inhale on the ENDS) by visual cues. In the second scan subjects received no cues, and were instructed to ‘smoke’ naturally, *ad libitum*.

## Methods

### Equipment

The ENDS devices were all widely commercially available at the time of testing. The five brands tested were: ‘Njoy’, ‘Puritane’, ‘Vype’, ‘Nucig’, and ‘Jasper & Jasper’. See table 1 for more details on each device. These were all first generation, or ‘cig-a-like’ devices, with a form factor designed to mimic traditional cigarettes. All were of similar construction, consisting of an outer body, a lithium-polymer (LiPo) battery, a pressure-activated LED, a resistive heating element and nicotine-soaked wadding. LiPo batteries are commonly used in devices designed for the MRI environment, as they contain no internal metal components. All were operated by inhalation, and all incorporated a LED at the end of the device, intended to mimic the burning ember of a traditional cigarette. We exploited this feature by using a custom-built opto-electronic device to record the output of the LED at the end of the ENDS (see supplementary methods for details). This enabled effective and easy monitoring of task compliance, and recorded a time-series to be used in the analysis of the naturalistic smoking task (see below).

**Table 1.** Characteristics of the ENDS devices used in the initial testing phase.

| Brand | Variety/model | Manufacturers stated nicotine content | Body construction | Length (mm) | Battery |
| --- | --- | --- | --- | --- | --- |
| Njoy | Gold | 3% (by weight) | Plastic | 84 | LiPo, 70mAh |
| Puritane | Disposable | 16mg/g | Metallic | 108 | LiPo, 240mAh |
| Vype | Bold | 18.6mg | Plastic | 85 | LiPo, 280mAh |
| Nucig | N/A | 18mg | Metallic | 106 | LiPo, 200mAh |
| Jasper & Jasper | Nano disposable | 16mg | Plastic; metal housing around heating element | 88 | LiPo, 170mAh |

**Table 2.**
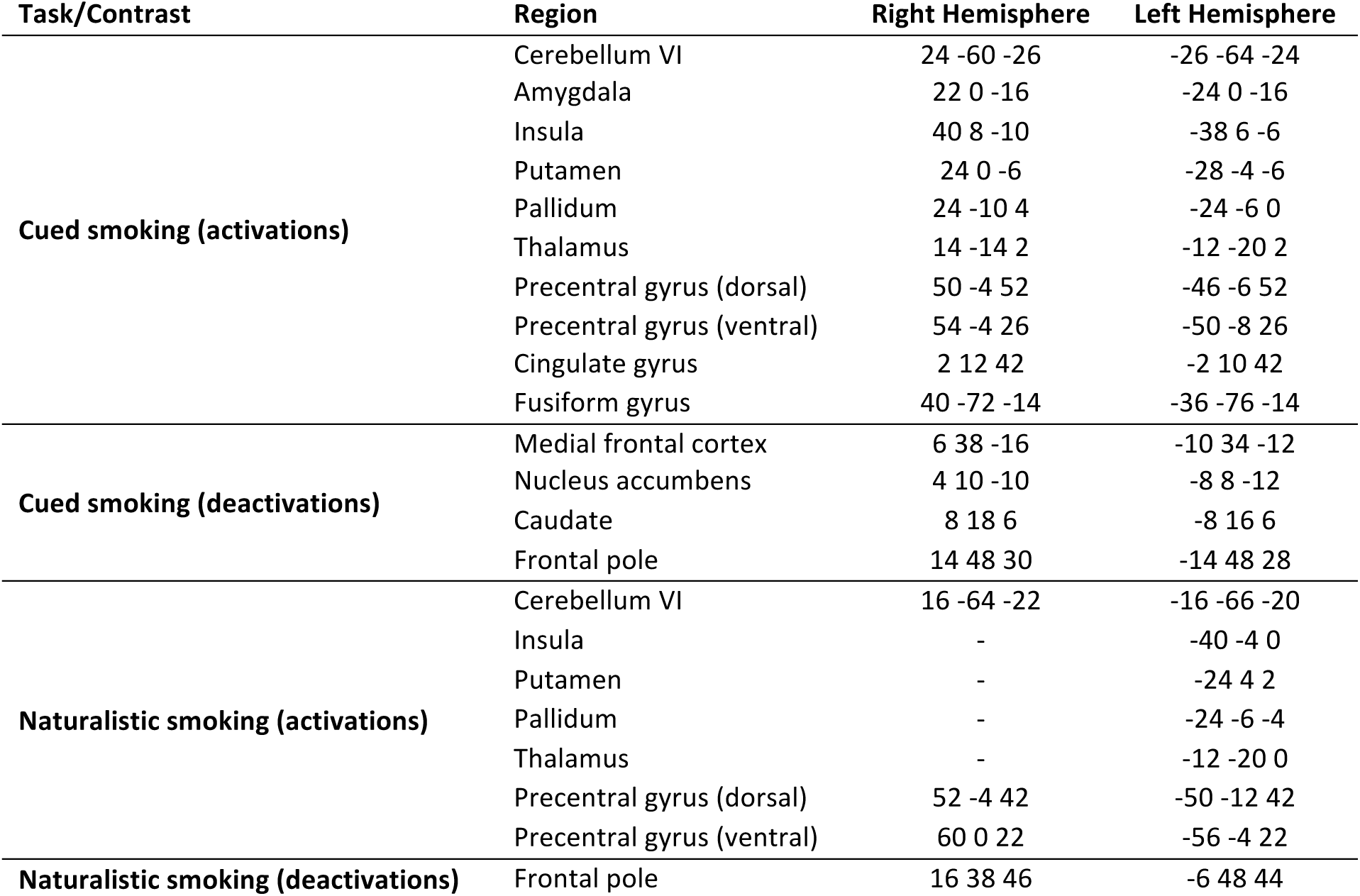
Coordinates of the approximate centre of activation clusters within anatomical regions for all experiments and contrasts. Coordinates are in MNI space.

### Initial Product Testing

We tested five different brands of commercially available ENDS for their suitability for use in the MRI environment. None were modified or tampered with in any manner. All five ENDS were immersed in a 2% agarose solution with 160mM NaCl for conductivity. This phantom was imaged on a Siemens 3T Verio MRI scanner (Siemens Healthcare, Erlangen Germany). Imaging consisted of a gradient-echo sequence; TR = 300 ms, TE 1 = 5.19 ms, TE 2 = 7.65 ms, flip angle = 60°, 1.5 x 1.5 x 2 mm voxels, 35 axial slices, bandwidth = 1520 Hz/pixel at two echo times: 1.93 and 4.39ms. A phase difference image between the two echo times was used to characterize the disturbance to the magnetic field (B0) from each ENDS. A spin-echo sequence was used to characterize disturbances to the applied RF (B1) field; TR = 5s, TE=8.5ms, 1.2 x 1.2 x 3.5mm voxels, 40 axial and coronal slices, bandwidth = 800Hz/pixel.

Subsequently, one subject (Author MBW) completed a scan using each of the five brands in a similar manner to the subjects in the main experiment (described below). These scans used a dual-echo, echo-planar imaging (EPI) sequence for BOLD contrast with 36 axial slices, aligned with the AC-PC axis (TR = 2000 ms, TE1 = 13 ms, TE2 = 31 ms flip angle = 80°, 3 mm isotropic voxels, parallel imaging factor = 2, bandwidth = 2298 Hz/pixel). The ENDS devices were therefore assessed for general magnetic susceptibility, and in particular for their potential to generate artefacts when used by a subject during a BOLD EPI acquisition.

### Subjects

Subjects were 11 (3 females) daily or semi-regular social smokers, who were in good general health. One (male) subject was subsequently excluded from analysis because of excessive (> 5 mm) head movement and failure to comply with the task, which left a final group of 10 (mean age of 29.1 years; SD = 5.91). The mean number of cigarettes smoked daily for this group was 10.1 (SD = 5.6). Subjects were not asked to abstain from smoking on the day of the scan, and were therefore not in a nicotine-deprived state.

### Tasks and scanning procedure

Data were acquired on a Siemens 3T Magnetom Trio MRI scanner (Siemens Healthcare, Erlangen, Germany), equipped with a 32-channel phased-array head coil. Subjects held the ENDS in their right-hand, with the right elbow cushioned so that they could comfortably hold it on their chest, close to their mouth, in order to minimise the hand movement required on each trial. Subjects could view a back-projected image on a screen in the rear of the scanner bore via a mirror mounted on the head-coil. They were instructed to try to avoid looking down (e.g. along the scanner bore, towards the ENDS) as they inhaled, in order to counteract the natural tendency to nod the head forward slightly when looking down.

Based on the results of the initial testing (see results section) the ‘Njoy’ device was selected as a good candidate device for use in the main experiment. A fresh ENDS was used for each subject, and the only modification made was the removal of the plastic end cap covering the LED. This was required in order to provide higher light output and a better signal for the opto-electronic recording device. The optical fibre was attached to the end of the ENDS by a rubber connector, and remained in place throughout the scan. The output of the device was recorded by a standard analogue-to-digital recording system (PowerLab 8/35, AD Instruments, Oxford UK), and physiological parameters (respiration via a respiratory belt around the subjects’ chest, and cardiac data via a pulse-oximeter on the index finger of the left hand) were recorded on the same system.

At the beginning of the scan session high-resolution T1-weighted anatomical images were acquired using a magnetization prepared rapid gradient echo (MPRAGE) sequence with parameters from the Alzheimer’s Disease Research Network (ADNI; 160 slices x 240 x 256, TR = 2300 ms, TE = 2.98 ms, flip angle = 9°, 1 mm isotropic voxels, bandwidth = 240Hz/pixel, parallel imaging factor = 2; Jack et al., 2008) along with B0 field-map images (sequence as described above). Subjects then completed two functional scans (sequence as described above) of ten minutes duration each. The first was the cued smoking task that consisted of 20 trials with inter-trial intervals that varied randomly between 20, 25, and 30 seconds (mean = 25s). On each trial, a three second countdown (3, 2, 1) was displayed in the centre of the screen, followed by the word ‘SMOKE’, displayed for two seconds. A fixation cross was present throughout the inter-trial interval. Participants were instructed to time their inhalations on the ENDS to coincide with the ‘SMOKE’ cue. The second task was the naturalistic smoking task, where there were no visual cues, and subjects were instructed to smoke ‘naturally’ throughout the ten minute scan.

### Data analysis

All analysis was conducted using FSL (Smith et al., 2004; Jenkinson et al., 2012) version 5.0. Preprocessing of the functional data involved removal of non-brain tissue, head-motion correction, spatial smoothing with a 6mm full-width-half-maximum Gaussian kernel, and high-pass temporal filtering with a cut-off of 100s. Additional correction for the effects of head-motion used the ICA-AROMA tool (ICA-based Automatic Removal Of Motion Artefacts; Pruim et al., 2015; Pruim, Mennes, Buitelaar, & Beckmann, 2015). For the cued smoking task data, first-level statistical (General Linear Model) models were created which contained a single regressor of interest. This time-series modelled two-second events defined by the occurrence of the smoking cue, convolved with a gamma function in order to produce a standard model of the haemodynamic response function. The first temporal derivative of this time-series was also included in the model. Statistical maps resulting from these analyses were coregistered to each subjects’ skull-stripped T1 anatomical image, and then to an image in standard stereotactic space (the MNI152 template provided with FSL). Analysis at the second (group) level computed a simple mean across subjects for the regressor of interest using FSL’s FLAME-1 model, and the results were thresholded at *Z* = 2.3 (*p* < 0.05, cluster-corrected for multiple comparisons).

For the naturalistic smoking task data, ‘smoking’ events were defined based on the recorded output of the opto-electronic device. This enabled a custom regressor of smoking events to be produced for each subject, using custom written MatLab (MathWorks Ltd.) code. The temporal derivative of the raw time-series was computed, and the peaks and troughs of the derivative time-series were used to define the start and end of each smoking event. In all other respects analysis of this task was identical to the cued smoking task.

Because of the established effects of physiological parameters on fMRI data (e.g. Birn et al., 2006; Harvey et al., 2008; Diukova et al., 2012), and because these tasks crucially depend on timed inhalations, analysis models of both sets of data also included physiological noise regressors. The recorded physiological (cardiac and respiratory) data was processed using the Physiological Noise Modelling (PNM) toolbox included with FSL (Brooks et al., 2008) and an additional 12 Fourier-expanded regressors were created to model cardiac and respiratory function. Additional analyses were conducted without the physiological noise regressors, in order to examine the effect of the noise modelling procedure.

## Results

### ENDS MR compatibility

None of the five ENDS contained ferromagnetic components such as might experience torque inside the magnetic field, however, two were found to have metallic bodies upon disassembly (see table 1). While no large torques would be expected, eddy currents will be induced in the conductive body from the MRI’s RF activity (Schenk, 1996). These eddy currents generate unwanted counter-acting magnetic fields as seen in supplementary figure 2, and may pose a safety risk from heating of the conductor. Further, most metals have a magnetic susceptibility, χ, far enough (> 10^−5^) from the χ of water that image distortion is expected. While an aluminium body would have a modest effect, nickel or stainless steel would create large image distortions. If the ENDS could be used parallel to the main magnetic field B0, the cylinder-shaped body would not perturb B0 outside the ENDS. However, when used normally the ENDS would be transverse to the main magnetic field; resulting in a dipole perturbation in cylindrical coordinates (ρ, φ) of 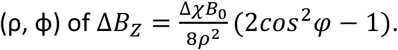. These characteristic dipole patterns are illustrated in supplementary figure 1.

**Figure 1.**
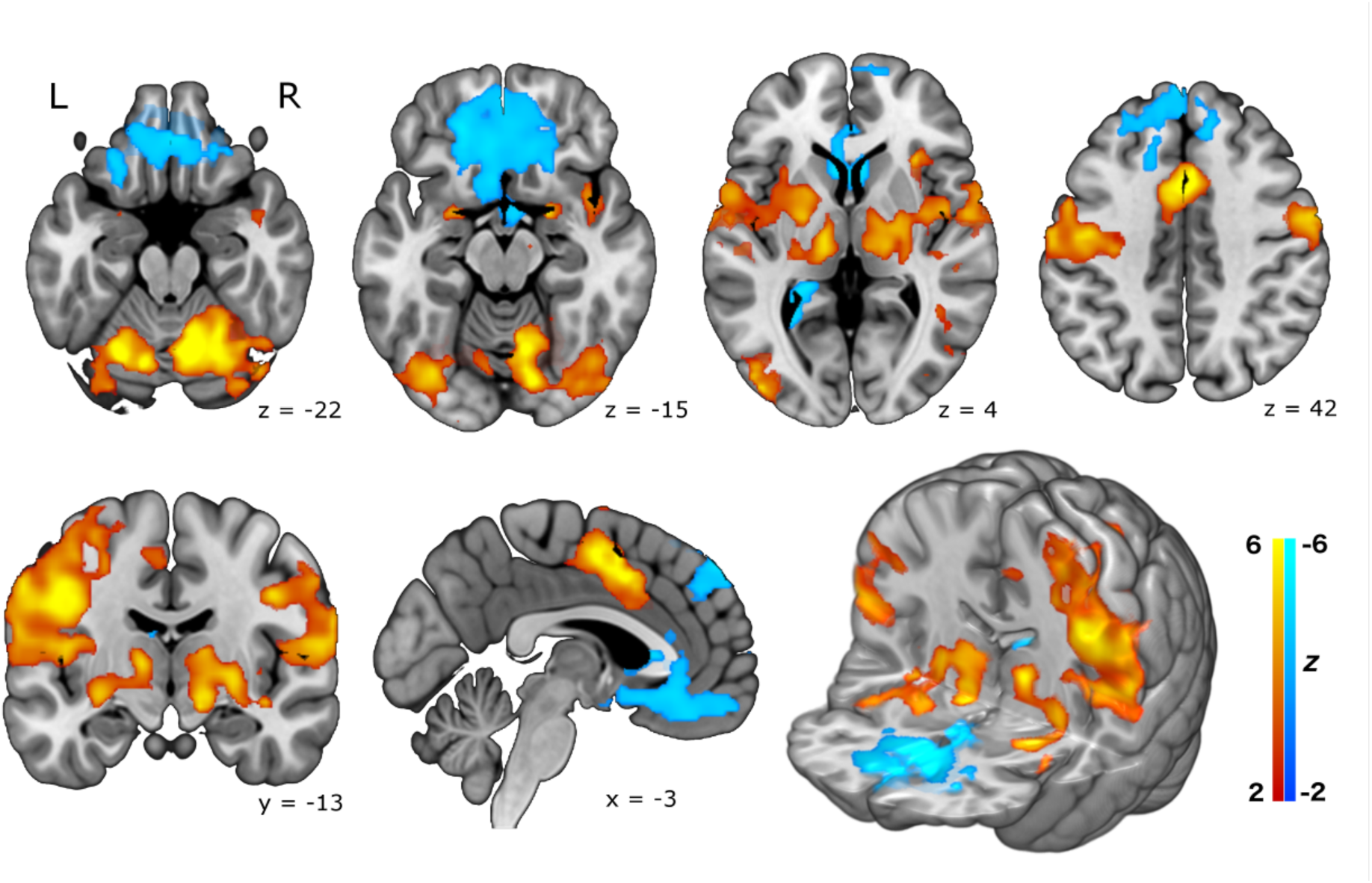
Brain activation in the cued smoking experiment, from a group mean analysis of 10 healthy smokers, including physiological noise modelling. Top row: axial slices. Bottom row: (from left) coronal slice, sagittal slice, and a 3D rendered image. Increased activity is seen in the amygdala, cerebellum, thalamus, putamen, globus pallidus, insula, cingulate gyrus and motor cortex. Reduced activity is seen in the ventral striatum, orbitofrontal cortex, and dorso-medial frontal regions. Statistical maps are shown thresholded at *z* > 2.3, *p* < 0.05 (cluster corrected for multiple comparisons) and neurological convention is used (L=Left, R=Right). Background anatomical image is a high-resolution version of the MNI152 T1 anatomical template.

**Figure 2.**
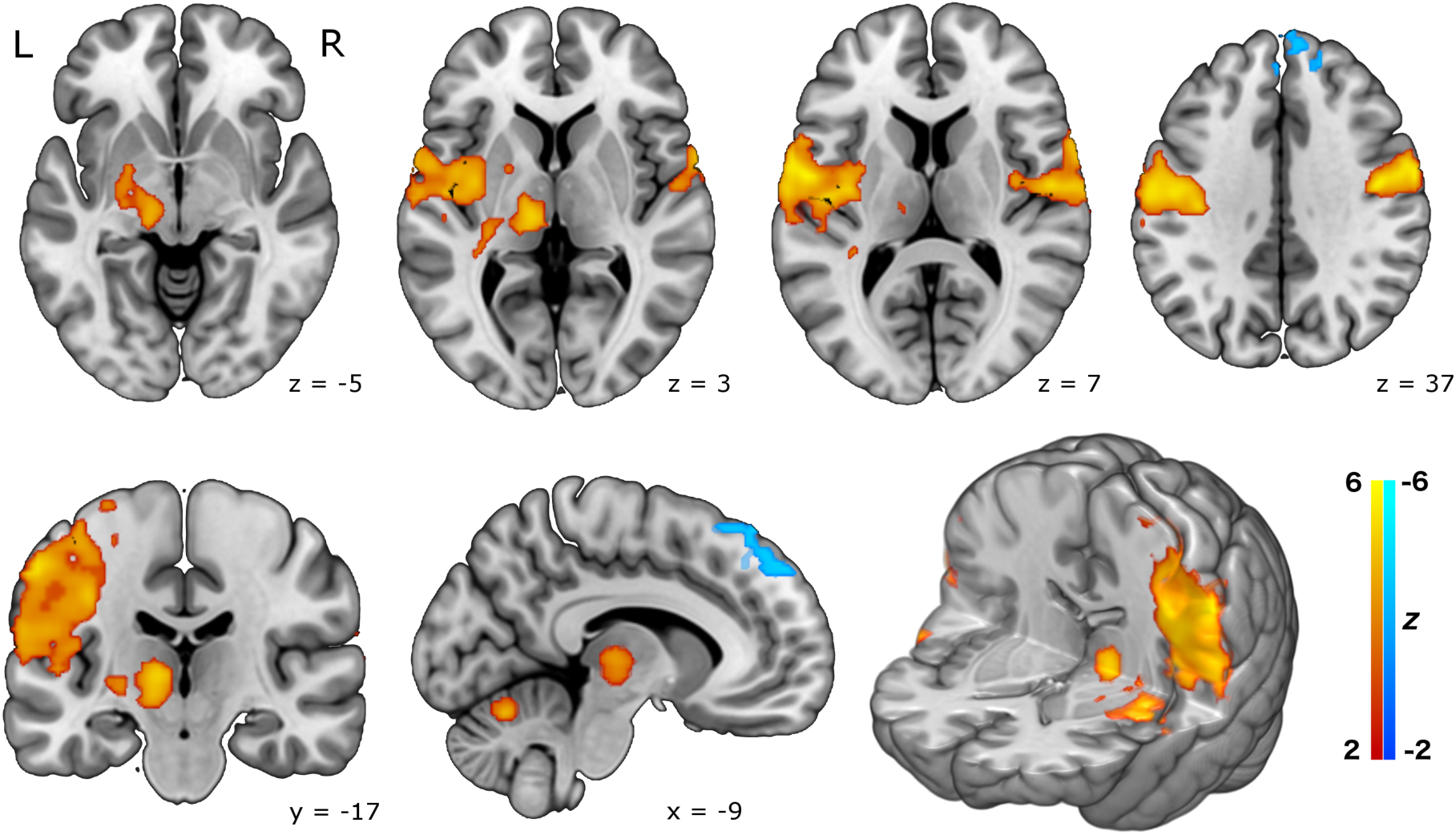
Brain activation in the naturalistic smoking experiment, from a group mean analysis of 10 healthy smokers, including physiological noise modelling. Top row: axial slices. Bottom row: (from left) coronal slice, sagittal slice, and a 3D rendered image. The pattern of activation is broadly similar to that seen in the cued experiment, but somewhat less robust, and less widespread. Statistical maps are shown thresholded at *z* > 2.3, *p* < 0.05 (cluster corrected for multiple comparisons) and neurological convention is used (L=Left, R=Right). Background anatomical image is a high-resolution version of the MNI152 T1 anatomical template.

The MRI environment had no apparent effect on the operation of any of the tested ENDS devices. The magnetic susceptibility artefacts produced by the different devices varied widely (see supplementary figure 2 for high and low-susceptibility examples). Unsurprisingly, the larger devices with metal construction produced the most detrimental effects, but even the smaller devices produced some areas of signal loss on the MRI phantom images. However, the smaller devices produced no obvious image artefacts in the test with a human subject, suggesting that they would be suitable for use in the main experiment.

### Task Results

In the main study, figure 1 illustrates the resulting pattern of brain activity that occurred in the cued-smoking task, time-locked to the smoking events. Large activation clusters are seen in the dorsal motor cortex in the left hemisphere, consistent with movements of the right hand during smoking trials. More lateral motor activity likely represents oro-facial movements on smoking trials. Other cortical regions strongly activated were the mid-insula, the amygdala, and the (dorsal) anterior cingulate gyrus. Strong activation of sub-cortical regions is also seen, principally in the thalamus, globus pallidus, and putamen. Relative deactivations associated with smoking events were also seen in a large region encompassing parts of the ventral striatum (nucleus accumbens and ventral caudate) and extending forward into orbitofrontal cortex. An additional area of deactivation was also seen in more dorsal frontal cortex.

Brain activity in the naturalistic smoking experiment showed a similar pattern, though less strongly, and less widespread, with significant clusters in motor cortex, thalamus, globus pallidus and putamen during smoking events. Relative deactivations were restricted to the dorsal frontal region in this analysis (see figure 2).

Additional analyses of both data-sets that did not include the physiological noise modelling regressors produced a highly similar set of results (see supplementary figure 3), suggesting that physiological effects are not a significant confound in the data.

## Discussion

These data demonstrate the feasibility of using ENDS in the MRI environment, and reveal the neural correlates of (simulated) smoking for the first time. A wide network of activated areas included cortical (motor cortex, insula, cingulate, amygdala) and sub-cortical (thalamus, putamen, globus pallidus, cerebellum) regions, with corresponding significant deactivations in ventral striatum and orbitofrontal cortex.

The most prominent result is the activation in left motor cortex which can plausibly be attributed to hand and oro-facial movements associated with the smoking task. The activation clusters in the right cerebellum are also most plausibly related to the motor components of the task. While certainly part of the behavioural repertoire of smoking, these results are unsurprising, and will not be discussed further. Of more interest are the results in other regions, particularly the thalamus and striatum. Previous work has identified differences in dopamine release in the caudate and putamen in smokers vs. non-smokers (Takahashi et al., 2008), and the striatum is widely considered to be a key set of brain structures that support addiction (Volkow et al., 2007; Robbins & Everitt, 2002; Everitt & Robbins, 2005), with interactions between nicotine and the mesolimbic dopamine system also well-established (for a review see Pierce & Kumaresan, 2006). Intriguingly, parts of the ventral striatum (nucleus accumbens, and ventral caudate) appear to show a relative signal *decrease* in response to smoking in this experiment, as well as a large region in the orbitofrontal cortex. One possible explanation for these regions of relative deactivation may be the shift from ventral to dorsal striatum that occurs as drug use becomes habitual and compulsive (Everitt & Robbins, 2013), and the finding that dysfunctional inhibitory mechanisms in the prefrontal cortex also play a role in addictive behaviour (Everitt & Robbins, 2013; Goldstein & Volkow, 2011). Janes et al. (2014) also recently showed that the connectivity of the orbitofrontal cortex was highly related to subjective craving measures. The current findings are consistent with the established role of the ventral striatum and frontal cortex in drug craving, and further suggest that activity in these areas is actually reduced during active drug consumption, with the rewarding aspect of consumption mediated by more dorsal striatal regions. The results are also highly consistent with the previous PET study by Berridge et al. (2010) which showed smoking-related euphoria was related to dopamine release in the dorsal, but not ventral, striatum. The general activation pattern involving the brain’s reward circuitry also implies that use of ENDS may have a similar rewarding effect as traditional cigarettes.

It is important to note that the current data say relatively little about the neural effects of nicotine. Although the ENDS used contained nicotine, the modelling of the brain response was time-locked to the action of smoking. Nicotine (absorbed into the blood through the vapour produced by the ENDS) is likely to only enter the brain between 5 and 20 seconds after each inhalation (Berridge et al., 2010), and modelling this effect within a conventional fMRI design would be difficult, since the timing of the nicotine ‘hit’ after each trial is somewhat uncertain. There is a large literature on nicotine and its pure pharmacological effects are relatively well understood (e.g. Stein et al., 1998; Yamamoto, Rohan & Goletiani, 2013). The current data stand as complementary to this literature, and provide a visualisation of brain processes related to the consumption of nicotine in a relatively naturalistic manner.

Results from the cued task were robust, while results from the naturalistic task were noticeably muted in comparison. There are several plausible explanations for this finding. Firstly in the naturalistic study the number of smoking events varied across subjects (mean = 28.8, SD = 15.4) and this produced more between-subject variance in the data. Secondly, the timing of events in the naturalistic study was determined by the subjects, and therefore did not necessarily conform to optimal principles of fMRI experimental design (Friston et al., 1999; Dale, 1999); this could affect the signal-detection power of the experiment. Thirdly, the naturalistic scan was always conducted after the cued scan and subjects were possibly nicotine-sated in the latter scan, leading to less activation in the striatal (i.e. more reward-related) regions. Fourth, the task demands of each scan were quite different, with one requiring active focus and attention on the external cue task, while the other did not. One other aspect of the design deserves comment, which is the use of a purely baseline control condition in both tasks. In this initial study we were concerned with demonstrating the technical feasibility of using ENDS in this way, but also with visualising the neural correlates of the entire behavioural and sensory repertoire of smoking. The comparison of ‘smoking’ with ‘not smoking’ (that is, a resting baseline) provides this. Future work may be concerned with dissecting the various components of the response using appropriate control conditions. For instance, sham-smoking with a non-functional device may be a useful control condition for subtracting out the motor aspects of the response.

Future work using similar methods may also be focussed on optimising the equipment, procedure, and analysis strategy. The market for electronic cigarettes has shown explosive growth in recent years, and new products are being launched almost on a daily basis. The first generation ‘cig-a-like’ devices used here are designed to mimic traditional cigarettes, and this visual mimicry may be helpful in reducing withdrawal symptoms (Dawkins et al., 2015). However the second or third generation devices currently on the market appear to be more effective at delivering nicotine, and thereby reducing craving and withdrawal symptoms (Lechner et al., 2015; Dawkins, Kimber, Puwanesarasa & Soar, 2015). Unfortunately these devices tend to be larger, heavier, and incorporate more metallic components, making them unsuitable for use in the MRI environment without substantial modification. The smaller, first generation devices (particularly those that have a plastic construction and use non-magnetic Lithium polymer or ‘LiPo’ batteries) are probably the best current option for this kind of work. The other useful feature of these devices is the end-mounted LED, which allows for easy monitoring of task compliance, with the simple device used here. Alternative analysis strategies for studies of this type may be focussed on visualising the trial-by-trial effect of nicotine on the brain, or the cumulative effect over the course of the experiment. This would be difficult because of the reasons mentioned above, but is perhaps feasible using a more flexible statistical model, or model-free analysis approaches such as Independent Components Analysis (ICA).

This demonstration of the feasibility of using ENDS in the MRI environment has served to validate an entirely novel approach to the study of cigarette dependence, and the more general brain mechanisms of addiction. We have also revealed for the first time the full neural effects of active (simulated) smoking, which includes activation in a network of cortical (motor, insula, cingulate, amygdala) and sub-cortical (putamen, thalamus) regions, with relative deactivation in ventral striatum and orbitofrontal cortex. Together with previous work on nicotine, and cue-reactivity in smokers, these findings provide a more complete picture of the neural effects associated with cigarette smoking, and addiction in general.

## Supplementary Material

### Methods

A custom-built opto-electronic device was used to record the light output of the LED of the ENDS during the scanning tasks. This consisted of a 10 metre optical fibre cable, attached to a small rubber connector that allowed it to be fitted to the ENDS, in the scanner room. The cable passed through a waveguide into the control room, and terminated in a light-tight box next to a photo-sensitive resistor. Wired in parallel with the resistor was a 1.5V power source (an AA battery) and a BNC cable. This circuit produced a voltage on the resistor whenever the ENDS was activated by an inhalation, and this time-series signal was recorded via the BNC connection by a standard analogue-to-digital data recorder (AD Instruments Powerlab). In this way compliance with the cued smoking task could be assessed, and the recorded data in the naturalistic smoking task could be used to define idiosyncratic ‘smoking’ events for each subject.

### Results

**Figure S1.**
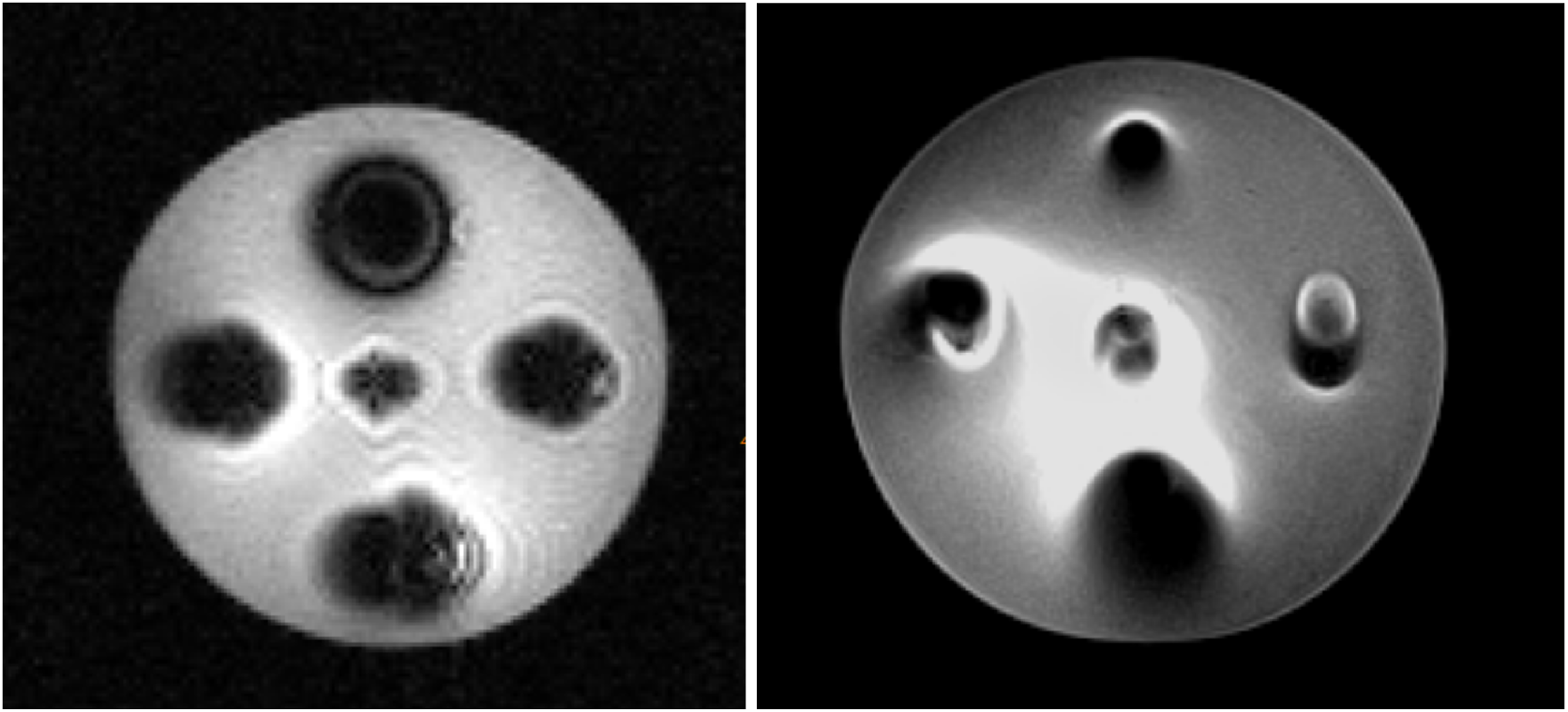
The five ENDS imaged with a gradient-echo sequence (left) shows characteristic dipole pattern disturbances from the different magnetic field susceptibilities found in the materials used to construct the varied ENDS. The same phantom images with a spin-echo sequence (right) removes the susceptibility-induced distortion and signal loss, but the RF field is compromised by the presence of conductors.

**Figure S2.**
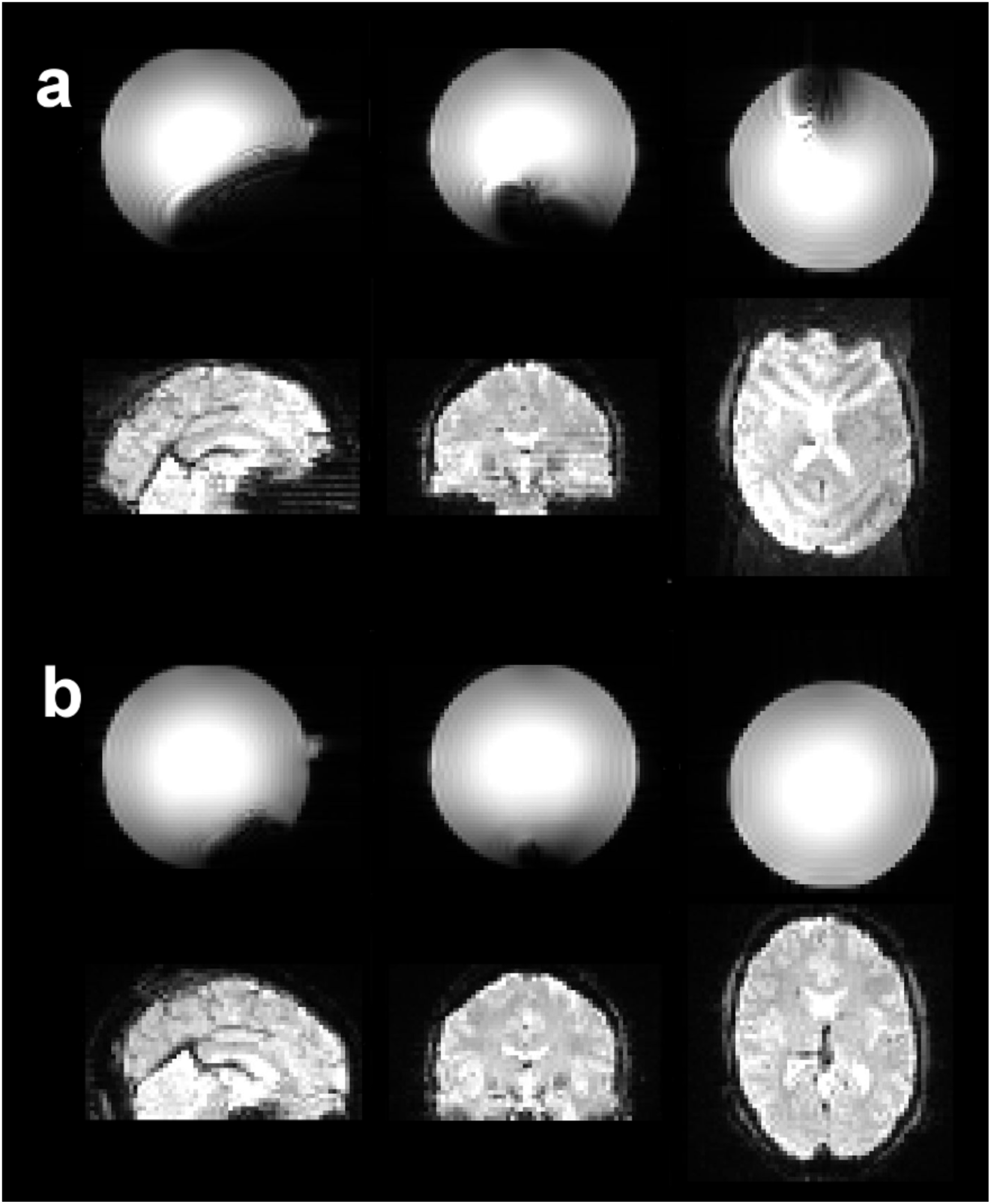
Results from initial testing of electronic cigarette products. a: An ENDS (‘Puritane’) with high magnetic susceptibility produces large regions of signal loss on a B0 magnitude image of an MRI phantom (top row). The same product also produces obvious artefacts on a BOLD EPI image when used by a human subject in the scanner (bottom row). b: A low magnetic susceptibility product (‘Njoy’) has much less effect on the B0 phantom image, and produces no obvious artefacts on BOLD EPI images from a human subject during use.

**Figure S3.**
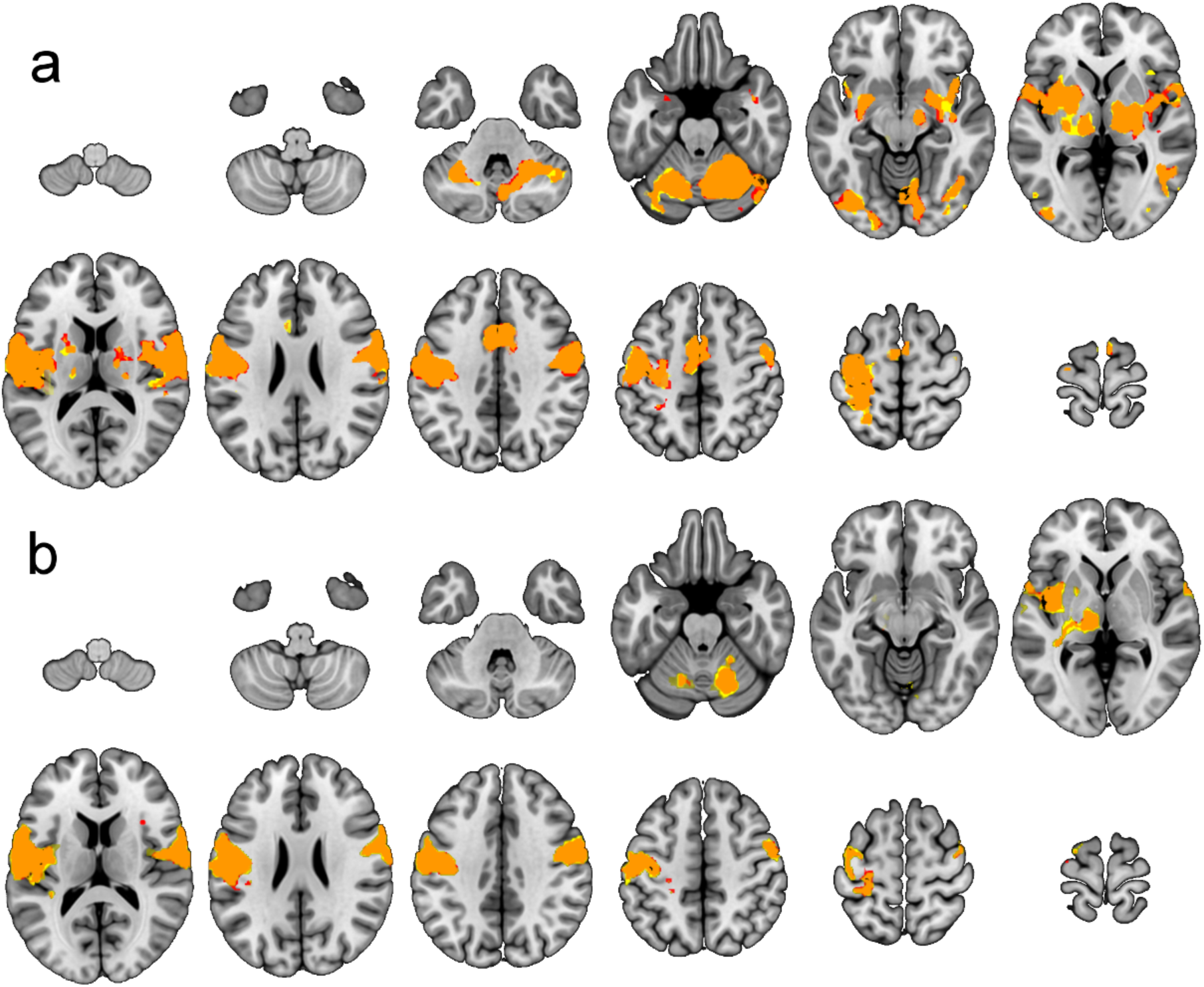
The effect of statistical modelling of physiological parameters (cardiac and respiratory effects) on the data from both experiments. Activation maps produced with no physiological modelling are shown in red, physiologically-corrected maps are shown in yellow, the overlap is in orange. a) Cued smoking experiment. b) Naturalistic smoking experiment. Effects of physiological noise modelling on the results are negligible, suggesting that these factors were not a significant confound in the data.

## Footnotes

1 One group has published two papers (Lindsey et al, 2009; Lindsey, Bracken & MacLean, 2013) validating a MRI compatible system to enable smoking during functional imaging, however to the current authors’ knowledge, they have never presented any fMRI data collected using the device.

## References

Balfour, D. (2004). The neurobiology of tobacco dependence: A preclinical perspective on the role of the dopamine projections to the nucleus. Nicotine & Tobacco Research, 6(6), 899–912. doi:10.1080/14622200412331324965

Berridge, M. S., Apana, S. M., Nagano, K. K., Berridge, C. E., Leisure, G. P., & Boswell, M. V. (2010). Smoking produces rapid rise of [11C]nicotine in human brain. Psychopharmacology, 209(4), 383–94. doi:10.1007/s00213-010-1809-8

Birn RM, Diamond JB, Smith MA, Bandettini PA. (2006) Separating respiratory-variation-related fluctuations from neuronal-activity-related fluctuations in fMRI. Neuroimage. 31:1536–1548.

Bullen, C., Howe, C., Laugesen, M., McRobbie, H., Parag, V., Williman, J., & Walker, N. (2013). Electronic cigarettes for smoking cessation: a randomised controlled trial. The Lancet, 382(9905), 1629–1637.

Brooks, J. C. W., Beckmann, C. F., Miller, K. L., Wise, R. G., Porro, C. A., Tracey, I., & Jenkinson, M. (2008). Physiological noise modelling for spinal functional magnetic resonance imaging studies. NeuroImage, 39(2), 680–692. doi:10.1016/j.neuroimage.2007.09.018

Cole, D. M., Beckmann, C. F., Long, C. J., Matthews, P. M., Durcan, M. J., & Beaver, J. D. (2010). Nicotine replacement in abstinent smokers improves cognitive withdrawal symptoms with modulation of resting brain network dynamics. NeuroImage, 52(2), 590–9. doi:10.1016/j.neuroimage.2010.04.251

Cosgrove, K. P., Wang, S., Kim, S. J., McGovern, E., Nabulsi, N., Gao, H.,… & Morris, E. D. (2014). Sex differences in the brain’s dopamine signature of cigarette smoking. Journal of Neuroscience, 34(50), 16851–16855.

Dale, A. M. (1999). Optimal experimental design for event-related fMRI. Human Brain Mapping, 8(2-3), 109–14. doi:10.1002/(SICI)1097-0193(1999)8:2/3<109::AID-HBM7>3.0.CO;2-W

David, S. P., Munafò, M. R., Johansen-Berg, H., Smith, S. M., Rogers, R. D., Matthews, P. M., & Walton, R. T. (2005). Ventral striatum/nucleus accumbens activation to smoking-related pictorial cues in smokers and nonsmokers: a functional magnetic resonance imaging study. Biological Psychiatry, 58(6), 488–94. doi:10.1016/j.biopsych.2005.04.028

Dawkins, L., Kimber, C., Puwanesarasa, Y., & Soar, K. (2015). First-versus second-generation electronic cigarettes: predictors of choice and effects on urge to smoke and withdrawal symptoms. Addiction, 110(4), 669–677.

Dawkins, L., Munafò, M., Christoforou, G., Olumegbon, N., & Soar, K. (2016). The effects of e-cigarette visual appearance on craving and withdrawal symptoms in abstinent smokers. Psychology of Addictive Behaviors, 30(1), 101.

Dawkins, L., Turner, J., Hasna, S., & Soar, K. (2012). The electronic-cigarette: effects on desire to smoke, withdrawal symptoms and cognition. Addictive behaviors, 37(8), 970–973.

Diukova, A., Ware, J., Smith, J. E., Evans, C. J., Murphy, K., Rogers, P. J., & Wise, R. G. (2012). Separating neural and vascular effects of caffeine using simultaneous EEG-FMRI: differential effects of caffeine on cognitive and sensorimotor brain responses. NeuroImage, 62(1), 239–49. doi:10.1016/j.neuroimage.2012.04.041

Domino, E. F., Ni, L., Domino, J. S., Yang, W., Evans, C., Guthrie, S.,… & Zubieta, J. K. (2013). Denicotinized versus average nicotine tobacco cigarette smoking differentially releases striatal dopamine. Nicotine & Tobacco Research, 15(1), 11–21.

Everitt, B. J., & Robbins, T. W. (2005). Neural systems of reinforcement for drug addiction: from actions to habits to compulsion. Nature neuroscience, 8(11), 1481–1489.

Everitt, B. J., & Robbins, T. W. (2013). From the ventral to the dorsal striatum: devolving views of their roles in drug addiction. Neuroscience & Biobehavioral Reviews, 37(9), 1946–1954.

Friston, K. J., Zarahn, E. O. R. N. A., Josephs, O., Henson, R. N. A., & Dale, A. M. (1999). Stochastic designs in event-related fMRI. Neuroimage, 10(5), 607–619.

Goldstein, R. Z., & Volkow, N. D. (2011). Dysfunction of the prefrontal cortex in addiction: neuroimaging findings and clinical implications. Nature Reviews Neuroscience, 12(11), 652–669.

Harvey, A. K., Pattinson, K. T. S., Brooks, J. C. W., Mayhew, S. D., Jenkinson, M., & Wise, R. G. (2008). Brainstem functional magnetic resonance imaging: disentangling signal from physiological noise. Journal of Magnetic Resonance Imaging, 28(6), 1337–44. doi:10.1002/jmri.21623

Health Act, 2006, Chapter 1. Available at: http://www.legislation.gov.uk/ukpga/2006/28/pdfs/ukpga_20060028_en.pdf (Accessed 22 November, 2016).

Jack, C. R., Bernstein, M. A., Fox, N. C., Thompson, P., Alexander, G., Harvey, D.,… & Dale, A. M. (2008). The Alzheimer’s disease neuroimaging initiative (ADNI): MRI methods. Journal of Magnetic Resonance Imaging,27(4), 685–691.

Janes, A. C., Farmer, S., Frederick, B. D., Nickerson, L. D., & Lukas, S. E. (2014). An increase in tobacco craving is associated with enhanced medial prefrontal cortex network coupling. PLoS ONE, 9(2), 1–5. doi:10.1371/journal.pone.0088228

Janes, A. C., Pizzagalli, D. A., Richardt, S., de B Frederick, B., Holmes, A. J., Sousa, J.,… & Kaufman, M. J. (2010). Neural substrates of attentional bias for smoking-related cues: an FMRI study. Neuropsychopharmacology, 35(12), 2339–2345.

Jenkinson, M., Beckmann, C. F., Behrens, T. E. J., Woolrich, M. W., & Smith, S. M. (2012). Fsl. NeuroImage, 62(2), 782–90. doi:10.1016/j.neuroimage.2011.09.015

Kalkhoran, S., & Glantz, S. A. (2016). E-cigarettes and smoking cessation in real-world and clinical settings: a systematic review and meta-analysis. The Lancet Respiratory Medicine, 4(2), 116–128.

Lechner, W. V., Meier, E., Wiener, J. L., Grant, D. M., Gilmore, J., Judah, M. R.,… & Wagener, T. L. (2015). The comparative efficacy of first-versus second-generation electronic cigarettes in reducing symptoms of nicotine withdrawal. Addiction, 110(5), 862–867.

Lindsey, K., Bracken, B., & MacLean, R. (2013). Nicotine content and abstinence state have different effects on subjective ratings of positive versus negative reinforcement from smoking. Pharmacology…, 103(4), 710–716. doi:10.1016/j.pbb.2012.11.012.Nicotine

Lindsey, K. P., Lukas, S. E., MacLean, R. R., Ryan, E. T., Reed, K. R., & Frederick, B. deB. (2009). Design and validation of an improved nonferrous smoking device for self-administration of smoked drugs with concurrent fMRI neuroimaging. Clinical EEG and Neuroscience : Official Journal of the EEG and Clinical Neuroscience Society (ENCS), 40(1), 21–30.

Nature Neuroscience (Editorial; 2014). Clearing the smoke. Nature Neuroscience, 17(8), 1013. doi:10.1038/nn.3777

Niaura, R., Shadel, W. G., Abrams, D. B., Monti, P. M., Rohsenow, D. J., & Sirota, A. (1998). Individual differences in cue reactivity among smokers trying to quit: effects of gender and cue type. Addictive behaviors, 23(2), 209–224.

Pierce, R. C., & Kumaresan, V. (2006). The mesolimbic dopamine system: The final common pathway for the reinforcing effect of drugs of abuse? Neuroscience and Biobehavioral Reviews, 30(2), 215–238. doi:10.1016/j.neubiorev.2005.04.016

Pruim, R. H. R., Mennes, M., van Rooij, D., Llera, A., Buitelaar, J. K., & Beckmann, C. F. (2015). ICA-AROMA: A robust ICA-based strategy for removing motion artifacts from fMRI data. NeuroImage, 112, 267–277. http://doi.org/10.1016/j.neuroimage.2015.02.064

Pruim, R. H. R., Mennes, M., Buitelaar, J. K., & Beckmann, C. F. (2015). Evaluation of ICA-AROMA and alternative strategies for motion artifact removal in resting state fMRI. NeuroImage, 112, 278–287. http://doi.org/10.1016/j.neuroimage.2015.02.063

Rahman, M. A., Hann, N., Wilson, A., Mnatzaganian, G., & Worrall-Carter, L. (2015). E-cigarettes and smoking cessation: evidence from a systematic review and meta-analysis. PloS one, 10(3), e0122544.

Robbins, T. W., & Everitt, B. J. (2002). Limbic-striatal memory systems and drug addiction. Neurobiology of learning and memory, 78(3), 625–636.

Robinson, T., & Berridge, K. (1993). The neural basis of drug craving: an incentive-sensitization theory of addiction. Brain Research Reviews, 8, 247–291.

Robinson, T. E., & Berridge, K. C. (2001). Incentive-sensitization and addiction. Addiction, 96(1), 103–14. doi:10.1080/09652140020016996

Rose, J. E. (2006). Nicotine and nonnicotine factors in cigarette addiction. Psychopharmacology, 184(3-4), 274–85. doi:10.1007/s00213-005-0250-x

Schenck, J. F. (1996). The role of magnetic susceptibility in magnetic resonance imaging: MRI magnetic compatibility of the first and second kinds. Medical physics, 23(6), 815–850.

Smith, S. M., Jenkinson, M., Woolrich, M. W., Beckmann, C. F., Behrens, T. E. J., Johansen-Berg, H.,… others. (2004). Advances in functional and structural MR image analysis and implementation as FSL. Neuroimage, 23, S208–S219.

Stein, E a, J Pankiewicz, H H Harsch, J K Cho, S a Fuller, R G Hoffmann, M Hawkins, S M Rao, P a Bandettini, and a S Bloom. 1998. “Nicotine-Induced Limbic Cortical Activation in the Human Brain: A Functional MRI Study.” The American Journal of Psychiatry 155 (8): 1009–15.

Takahashi, Hidehiko, Yota Fujimura, Mika Hayashi, Harumasa Takano, Motoichiro Kato, Yoshiro Okubo, Iwao Kanno, Hiroshi Ito, and Tetsuya Suhara. 2008. “Enhanced Dopamine Release by Nicotine in Cigarette Smokers: A Double-Blind, Randomized, Placebo-Controlled Pilot Study.” The International Journal of Neuropsychopharmacology 11 (3): 413–17. doi:10.1017/S1461145707008103.

Volkow, N. D., Fowler, J. S., Wang, G. J., Swanson, J. M., & Telang, F. (2007). Dopamine in drug abuse and addiction: results of imaging studies and treatment implications. Archives of neurology, 64(11), 1575–1579.

Yamamoto, R. T., Rohan, M. L., Goletiani, N., Olson, D., Peltier, M., Renshaw, P. F., & Mello, N. K. (2013). Nicotine related brain activity: The influence of smoking history and blood nicotine levels, an exploratory study. Drug and Alcohol Dependence, 129(1-2), 137–144. doi:10.1016/j.drugalcdep.2012.10.002

